# Development of radial frequency pattern perception in macaque monkeys

**DOI:** 10.1101/2024.02.21.581393

**Authors:** C. L. Rodríguez Deliz, Gerick M. Lee, Brittany N. Bushnell, Najib J. Majaj, J. Anthony Movshon, Lynne Kiorpes

## Abstract

Infant primates see poorly, and most perceptual functions mature steadily beyond early infancy. Behavioral studies on human and macaque infants show that global form perception, as measured by the ability to integrate contour information into a coherent percept, improves dramatically throughout the first several years after birth. However, it is unknown when sensitivity to curvature and shape emerges in early life. We studied the development of shape sensitivity in eighteen macaques, aged 2 months to 10 years. Using radial frequency stimuli (RFS), circular targets whose radii are modulated sinusoidally, we tested monkeys’ ability to discriminate RFS from circles as a function of the depth and frequency of sinusoidal modulation. We implemented a new 4-choice oddity task and compared the resulting data with that from a traditional 2-alternative task. Behavioral performance at all radial frequencies improved with age. Performance was better for higher radial frequencies, suggesting the developing visual system prioritizes processing of fine visual details that are ecologically relevant. By utilizing two complementary methods, we were able to capture a comprehensive developmental trajectory for shape perception.

## Introduction

Much research has been undertaken over the last 50 years to address the questions of what infants can see and when vision matures to adult levels. Although newborns can resolve and discriminate simple contrasting patterns that are matched for total area, luminance, number of elements and contour lengths (Fantz, 1963, 1964; Hershenson, 1964), their ability to resolve fine detail is very poor (Teller, 1981). Over the succeeding weeks, months and years, fine spatial resolution and many other aspects of form vision become fully developed. Importantly, distinct features of visual function emerge at different ages, and thereafter they improve at different rates throughout early infancy and childhood (see (Teller & Movshon, 1986; Teller, 1997; Johnson, 2011; Kovacs et al., 1999; Braddick & Atkinson, 2011; Atkinson & Braddick, 2020), for reviews). Perceiving a visual object is a hierarchical process, which builds upon the processing of local cues such as oriented edges, angles and curves. By integrating information at different spatial scales and levels of complexity, the brain can generate coherent, global representations of form (Kimchi, 1992; Kimchi et al., 2005). Existing literature on the development of global perception in children is inconclusive. Some reports indicate that young infants can distinguish global forms (Gerhardstein et al., 2004; Nayar et al., 2015), but other work using a variety of stimuli suggests that infants’ ability to integrate spatial cues in service of global form perception is poor and continues to improve well into childhood (Schwartz, Day, & Cohen, 1979; Kimchi, Hadad, et al., 2005; Scherf et al., 2009; Hadad, Maurer, & Lewis, 2010). For example, the ability to link segments of lines across space (contour completion), is measurable by 3 months of age on tasks using Gabor patches but continues to improve well into adolescence (Kovács, 1996; Kovacs, Kozma, et al., 1999; Gerhardstein, Kovacs, et al., 2004). Glass patterns are a canonical global form stimulus, consisting of fields of dot pairs that can be oriented and arranged to elicit the perception of global structure (Glass, 1969; Glass & Pérez, 1973). Braddick et al. (2000) identified weak sensitivity to Glass patterns by 4-months of age. Subsequent studies found that behavioral performance on Glass pattern discrimination tasks improves steadily throughout the first decade of life (Lewis et al., 2004).

Another example of global form perception is the perception of illusory contours. Adults can reliably distinguish objects that are partially occluded or that are not defined by physical boundaries, while it is unclear at what age young children can reliably do so. Performance does not reach adult levels of discrimination until late childhood or adolescence (Nayar, Franchak, et al., 2015). A big limitation of human studies is that longitudinal data are rarely available and typically focus on one or a few selected ages, leaving large age gaps and providing us with only snapshots of performance at particular points in developmental time. Moreover, only limited amounts of data can be obtained from a given subject in most cases. These limitations can be overcome by studying animal models. Macaque monkeys are particularly well-suited for this work because the monkey visual system is highly similar to that of humans, but visual performance develops roughly 4 times faster than humans, making it possible to track changes in visual behavior throughout development (Teller & Movshon, 1986; Teller, 1997). The development of the macaque visual system has been extensively characterized by our lab and others, both at a behavioral and neurophysiological level (see, (Boothe, Dobson, & Teller, 1985; Blakemore & Vital-Durand, 1986; Kiorpes & Kiper, 1996; Teller, 1997; Chino, Smith III, et al., 1997; Kiorpes & Movshon, 2004; Daw, 2006; Kiorpes & Movshon, 2014; Danka Mohammed & Khalil, 2020). Prior studies of the development of global form perception in monkeys, as measured using oriented Gabor patches in a contour integration task and Glass patterns, showed that sensitivity is measurable around 16 weeks. These abilities continue to improve into the second year of postnatal life (Kiorpes & Bassin, 2003; Kiorpes et al., 2012). Perception of simple textures, however, is already evident by 6 weeks of age, and matures relatively early (El-Shamayleh, Movshon, & Kiorpes, 2010). The common denominator shared by all of these types of global form stimuli is the underlying requirement of detecting and integrating local contrast, orientation and spatial frequency cues across space while discriminating fine details in the image (Wilson, Wilkinson, & Asaad, 1997; Wilkinson, Wilson, & Habak, 1998; Badcock, Clifford, & Khuu, 2005; Wang & Hess, 2005; Badcock & Clifford, 2006; Poirier & Wilson, 2006).

In this work, we assessed infant macaques’ sensitivity to radial deformation of circular contours using radial frequency stimuli (Wilkinson, Wilson, & Habak, 1998). Radial frequency patterns are a useful stimulus class to assess global shape processing because these patterns represent a wide array of smooth closed shapes that are commonly seen in nature, like fruits and flowers (Wilkinson, Wilson, & Habak, 1998; Hess, Achtman, & Wang, 2001; Hess, Wang, & Dakin, 1999; Jeffrey, Wang, & Birch, 2002; Loffler, Wilson, & Wilkinson, 2003). Perception of these patterns requires integration of information across space to accurately perceive the overall shape of the stimulus. Evidence shows that even though the circular contour is continuous, detecting deformations from circularity relies on comparing orientation information at different points along the contour rather than depending on local cues at a single point (Wilkinson, Wilson, & Habak, 1998; Hess, Wang, & Dakin, 1999; Loffler, Wilson, & Wilkinson, 2003; Jeffrey, Wang, & Birch, 2002). Adults show remarkably high sensitivity to radial frequency patterns, with thresholds reaching hyperacuity levels (Wilkinson, Wilson, & Habak, 1998). In other words, adult observers are able to resolve spatial distinctions at a scale finer than simple spatial resolution (Westheimer, 2010). Previous hyperacuity studies in infant humans and monkeys using traditional Vernier stimuli showed a protracted developmental time course for this ability as compared to grating acuity (Manny & Klein, 1984; Shimojo et al., 1984; Shimojo & Held, 1987; Norcia, Manny, & Wesemann, 1988; Kiorpes, 1992; Zanker et al., 1992; Brown, 1997; Carkeet, Levi, & Manny, 1997; Skoczenski & Norcia, 2002; Wang et al., 2009; Kiorpes, 2015), suggesting that radial frequency discrimination might also show protracted development. One previous study of young human infants showed that they displayed poor sensitivity to radial frequency patterns, with a period of rapid maturation between 4-6 months of age followed by an extended period of slower improvement (Birch, Swanson, & Wang, 2000). The developmental trajectory of sensitivity to radial deformations in macaques has not yet been studied. We tested macaque monkeys (aged 2 months to 10 years) on radial frequency discrimination tasks. We describe the development of global form sensitivity within and across subjects, demonstrating that even the youngest monkeys in our cohort could discriminate between radial frequency patterns and circles, but their behavioral sensitivity continued to improve well beyond the first year of life. We found it expedient to use a novel 4-choice oddity task to track performance in younger animals, which complemented data obtained in older animals using traditional forced-choice methods. We show that both tasks reliably capture the developmental trajectory of global form perception, with a predictable difference in sensitivity between tasks. Overall, our data reflect a developmental improvement in the perception of global shape information that is contingent on the spatial extent over which the visual system is able to integrate signals.

## Methods

### Subjects

Eighteen visually normal pig-tailed macaque monkeys (*Macaca nemestrina*) participated in this study, 4 females, 14 males. At the time of testing, their ages ranged from 8 to 491 weeks. Twelve subjects were tested at multiple age points throughout development and six were tested cross-sectionally. All animal care and experimental procedures were approved by the Institutional Animal Care and Use Committee of New York University and were in compliance with the guidelines established in the NIH Guide for the Care and Use of Laboratory Animals.

**Table 1:** Behavioral subjects. Identification, ages at which at least 1 radial frequency type was tested, task modality are listed in the first three columns. Single asterisk indicates ages tested with the preferential looking paradigm, double asterisks indicates ages tested with the 4-choice oddity task for subjects that were also tested on the 2-AFC task. The last two columns indicate whether the subject was tested with both eyes (BINO) or one eye at a time (MONO), and whether they were tested using a corrective lens (R / L). Only data from the dominant eye was included for these subjects.

| Subject | Ages tested (wks) | Task | Bino/Mono | Lens (R / L) |
| --- | --- | --- | --- | --- |
| M1 (female) | 16, 20, 60, 80, 88, 92 | 4-choice | BINO | – / – |
| M2 (female) | 16, 20, 56, 84, 88 | 4-choice | BINO | – / – |
| M3 | 12, 16, 28, 32, 52 | 4-choice | BINO | – / – |
| M4 (female) | 28, 32 | 4-choice | BINO | – / – |
| M5 | 16, 24, 28, 32, 48, 60 | 4-choice | BINO | – / – |
| M18 | 8, 12, 24, 32 | 4-choice | BINO | – / – |
| M6 | 68, 180** | Both | BINO | – / – |
| M7 | 67, 180** | Both | BINO | – / – |
| M8 (female) | 20*, 77, 181 | 2AFC | BINO | – / – |
| M9 | 17*, 73, 173 | 2AFC | BINO | – / – |
| M10 | 30 | 2AFC | BINO | – / – |
| M11 | 52, 158 | 2AFC | BINO | – / – |
| M12 | 48, 135, 199, 230 | 2AFC | MONO | +1.0D / +1.0D |
| M13 | 353 | 2AFC | MONO | – / + 1.75D |
| M14 | 125, 237 | 2AFC | BINO | – / – |
| M15 | 99 | 2AFC | BINO | – / – |
| M16 | 491 | 2AFC | MONO | +2.0D / +2.25D |
| M17 | 433 | 2AFC | MONO | +1.25D / +1.0D |

### Visual Stimulus Display

Stimuli were generated using custom software developed in our lab. The display consisted of a gamma-calibrated CRT monitor, placed at a distance determined by each behavioral set up.

For the 2-AFC experiments, stimuli were presented on a monitor with a resolution of 1024×768 at 100Hz and a mean luminance of 30 cd/m^2^. The screen subtended 43.6 degrees at a viewing distance of 50 cm and pixel size was 0.043 degrees in the reinforced-looking paradigm. Older animals tested on a 2AFC bar-pulling paradigm worked at a viewing distance of 100 cm, where the screen subtended 22.6 degrees and pixel size was 0.022 degrees. The display for the 4-choice experiments was a monitor with a resolution of 1280×960 at 120 Hz and a mean luminance of 28 cd/m^2^, combined with an EyeLink 1000 (SR Research) eye tracker. Viewing distance was 114 cm, at which the monitor subtended 19.9 degrees and pixel size was 0.016 degrees. Stimulus size was equated across the different viewing set ups, independently of pixel size.

### Radial Frequency Stimuli

To characterize the development of global form sensitivity, we used radial frequency stimuli (Figure 1A), a family of circular contours in which the cross-section of the outline’s luminance profile is determined by a fourth-derivative Gaussian. These were designed by Wilkinson, Wilson, and Habak (1998) with the goal of defining shape space parametrically in order to measure shape selectivity and sensitivity in psychophysical experiments. The control–distractor–stimulus in our study is the base circle. Unique stimuli can be generated from the base circle by sinusoidally modulating the contour a given number of times to create different shapes. The amplitudes of these modulations can be parametrically adjusted so that the stimulus is either like the base circle or not (Figure 1B). For our study, we kept the stimulus radius fixed at 1.5 degrees and the carrier spatial frequency at 2 cycles per degree, which is typically peak contrast sensitivity for animals across this age range.

**Figure 1:**
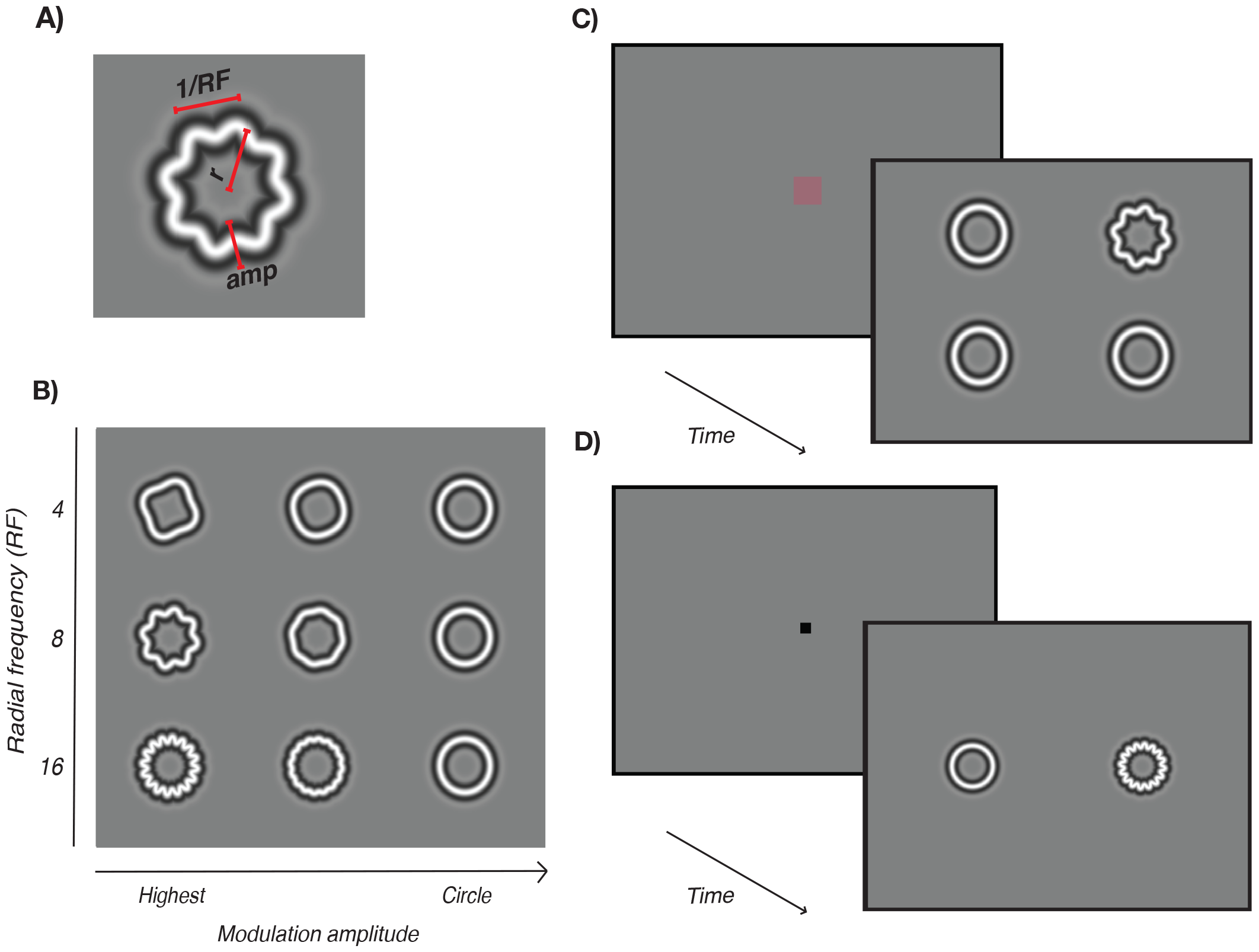
Radial Frequency Stimuli and task diagrams. A) Each stimulus is defined by a radial frequency (RF), amplitude (amp), and size (mean radius, r). B) We used the three radial frequency types (4, 8, 16), shown on the ordinate. Modulation amplitude is plotted along the abscissa, depicting a sample of modulations that are increasingly harder to discriminate from the base circle. C) On each trial of the 4-choice oddity task, subjects were required to fixate on the central fixation point. Four stimuli then appeared on the screen, equidistant from the center. Three stimuli were distractor circles, the target was a radial frequency pattern of varying amplitude modulation. Subjects are required to identify and fixate on the target for 400 ms. D) On each trial of the 2-AFC task, 2 stimuli appear on the screen, one is a distractor circle and the target is a radial frequency pattern. Subjects are required to indicate the side on which the radial frequency pattern appeared.

### Psychophysical experiments

#### 4-choice oddity task

We designed a new 4-choice oddity discrimination task that implements high-speed eye tracking (EyeLink 1000, SR Research), task automation (MWorks) and reward systems to reduce the time needed for training and testing subjects. This method allowed us to assess sensitivity to radial frequency stimuli across development starting at young ages, with an automated procedure, while maximizing the number of trials a subject completes in each session. Prior to testing, we trained five infant monkeys to sit in a custom-designed testing chair and fixate on a screen, head-free, starting at∼6 postnatal weeks. Two juveniles that had previously been tested using 2-AFC (described below) were also trained to perform the four-alternative oddity discrimination task. A total of seven subjects were tested on this paradigm.

A trial began with a gray screen and a 1-2 degree red, central fixation square. A trial was initiated when the subject fixated on the center of the screen, which triggered the disappearance of the fixation square, followed by a 200 ms delay and the subsequent presentation of a 2 by 2 array of stimuli of equal size (3 degrees) centered at ±3.2 degrees in both directions such that each stimulus was ∼4.5 degrees eccentric from the center of the screen (Figure 1C). Three of the stimuli were distractors (circles) and one was the target, a modulated radial frequency stimulus. Subjects were required to saccade toward the target and maintain fixation on it for 400 ms within a 5-degree window for the trial to be considered a hit and rewarded with age-appropriate liquids. A small proportion (∼5%) of catch trials were included to assess for any positional bias. To ensure accurate classification of trials, we extracted saccade and eye position data from MWorks output files. With this, we classified trials based on eye position and the subject’s behavior. Correct trials were any trial during which the animal fixated on the target for the full 400 ms. Failed trials were parsed into two categories: a miss if the animal fixated a distractor, and ignored/no-decision trials, in which the animal did not fixate any location for the full 400 ms. Ignored trials were excluded from our analysis. We randomly chose difficulty on a trial to trial basis, using the method of constant stimuli. To encourage cooperative behavior, it was our standard procedure to allow a larger amount of easy trials throughout the session. To characterize their perceptual thresholds, data were collected using a method of constant stimuli with a range of octave-spaced modulation amplitudes for a three radial frequency types (4, 8, 16).

#### 2-alternative forced choice task

We used standard operant conditioning methods to test visual performance in as previously described ((Boothe et al., 1988; Kiorpes & Movshon, 1989; Kiorpes, Kiper, & Movshon, 1993; Kozma & Kiorpes, 2003; El-Shamayleh, Movshon, & Kiorpes, 2010). Fourteen monkeys were trained to perform an operant two-alternative forced-choice discrimination task. At the time of testing, their ages ranged from 17 to 491 weeks, allowing us to track development of form sensitivity over the long term. Infants were allowed to roam freely in a cage equipped with a face mask. Photocells embedded in the mask signaled when they placed their faces in the mask, and triggered a new trial.

Stimuli appeared 5 degrees from the center on either side of the central fixation point. The stimulus was present on the screen for a minimum of 500 ms and a maximum of 3000 ms before the trial ended. The animals were shown a modulated radial frequency stimulus on one side and a circle on the other. Animals were trained on a reinforced looking paradigm (Kiorpes & Kiper, 1996), generating a saccade to the target (for macaques younger than 20 weeks) or pulling a bar to indicate whether the target stimulus was located on the left or the right side of the screen (Figure 1D). Monkeys received a small juice reward for correct trials and an error tone followed by a short pause if incorrect.

The goal was to find a range of 5 modulation amplitudes spanning the subject’s perceptual threshold from near perfect (100%) to near chance (25% on the 4-choice oddity and 50% on the 2-AFC). Difficulty was increased by parametrically reducing the modulation amplitude of the pattern, making it harder for the subject to discriminate between circles and radial frequency stimuli. Once threshold values were identified, we then counterbalanced by collecting a final session per radial frequency type in the reverse order as they were initially presented, to control for testing and order effects.

### Data Analyses

Wilkinson, Wilson, and Habak (1998) reported that, for human observers, sensitivities were typically better for higher radial frequencies, which may reflect differences in curvature-processing mechanisms across radial frequency types. Therefore, we measured modulation thresholds separately for each RF type. Sessions collected within the same 7-day span were combined for a given age point.

#### Threshold estimation

For the 2-AFC task, thresholds were estimated based on 75 trials per stimulus condition. Threshold values for this task were obtained by using a probit regression to determine the interpolated value corresponding to 75% correct performance. To compare the developmental trajectories derived from our oddity task and data collected using our standard 2-AFC task (e.g. (Kiorpes, Price, et al., 2012)), we matched our 4-choice threshold distributions by identifying the performance criteria that corresponds to 75% correct performance on a 2-AFC task, at which d’ = 0.95 (Hacker & Ratcliff, 1979).

Thresholds for the 4-choice oddity task were estimated on a minimum of 100 trials per stimulus condition to ensure robust definition of slope and threshold values. We fit a cumulative of the Weibull distribution to the performance data to estimate the behavioral threshold (Wichmann and Hill, 2001a). We used a maximum likelihood fitting process to extract a shared slope and lapse parameter, and a separate threshold for each individual measurement. Thresholds were computed as the intercept of the Weibull at 53.7% correct, corresponding to a d’ = 0.95. We estimated variability using a parametric bootstrap by fixing slope and lapse rates to the values extracted from the original fitting routine.

#### Sensitivity measurements

Developmental trends for all tasks were defined by fitting a Michaelis-Menten function to fitted sensitivity (1/threshold) data as a function of age using the equation:

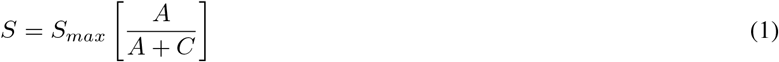

Where S is the fit sensitivity and *S*_*max*_ is the peak sensitivity, *A* is the subjects’ age in weeks, and *C* is the criterion age at which sensitivity reached half its maximum value (semi-saturation point). Fits were computed by minimizing the squared error of the model predictions. The data were bootstrapped 1000 times by re-sampling data for each radial frequency with replacement to extract parameter *C* (half-max age) on each iteration and compute 95% confidence intervals.

## Results

### Measuring radial frequency sensitivity

For each of our animals, we measured perceptual thresholds for radial frequency stimuli in a series of 4-choice oddity and/or 2-AFC experiments. Data were collected with a combination of longitudinal and cross-sectional testing. We used a block design to test each radial frequency (RF4, RF8 or RF16) separately. We examined performance on catch trials, in which all stimuli were identical, to evaluate potential biases toward particular screen locations. No systematic positional response biases were evident across subjects based on catch trial accuracy.

Figure 2 shows example psychometric curves measured for two infant macaques tested on the 4-choice task at 16 weeks of age. Performance at a given age varied across the radial frequencies tested, as shown by the horizontal separation between the red (RF4), blue (RF8) and green (RF16) curves. For all 3 radial frequencies tested, our new 4-choice oddity task yielded lawful psychometric functions. In both cases, sensitivity to modulation for the highest RF was better. We took the threshold for each function to be the amplitude supporting 53.7% correct performance in this task (indicated by arrows).

**Figure 2:**
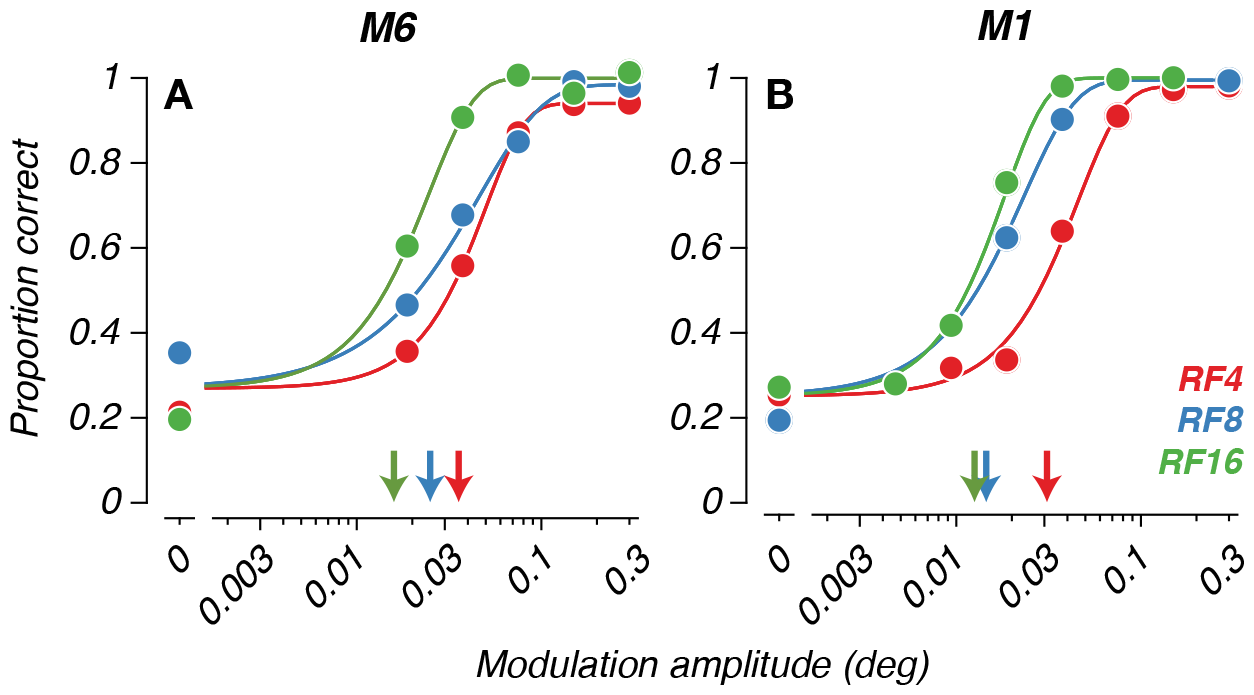
Example 4-choice data from two 16-week-old macaques. (A and B) Psychometric curves computed for infant macaques tested on each of three radial frequencies. Mean performance (solid circles) for each modulation amplitude level was fit with a Weibull. Radial frequency type is indicated by color (red = RF4; blue = RF8; green = RF16). Isolated points along the ordinate indicate performance on catch trials. Arrows indicate threshold amplitudes.

We tested a total of 18 animals, 13 at more than one age. Two (M6 and M7) were tested on both paradigms at different ages. Figure 3 shows radial frequency discrimination performance as sensitivity (inverse of threshold) to modulation amplitude, as a function of radial frequency for those 13 animals. Test age in weeks is indicated next to each data set. In most cases, sensitivity improved with age. However, the data also reveal some individual variability in developmental trajectories across subjects. In some cases, like panels B-D, for example, the animal’s motivation may have fluctuated over sessions and age, resulting in lower estimated sensitivities than one would expect given the animal’s earlier performances. Overall, the developing improvement in modulation sensitivity is clear in spite of some variability, possibly reflecting maturation of neural mechanisms for radial frequency processing. Moreover, the pattern of higher modulation sensitivity for higher RFs was evident across most subjects, and for each task modality.

**Figure 3:**
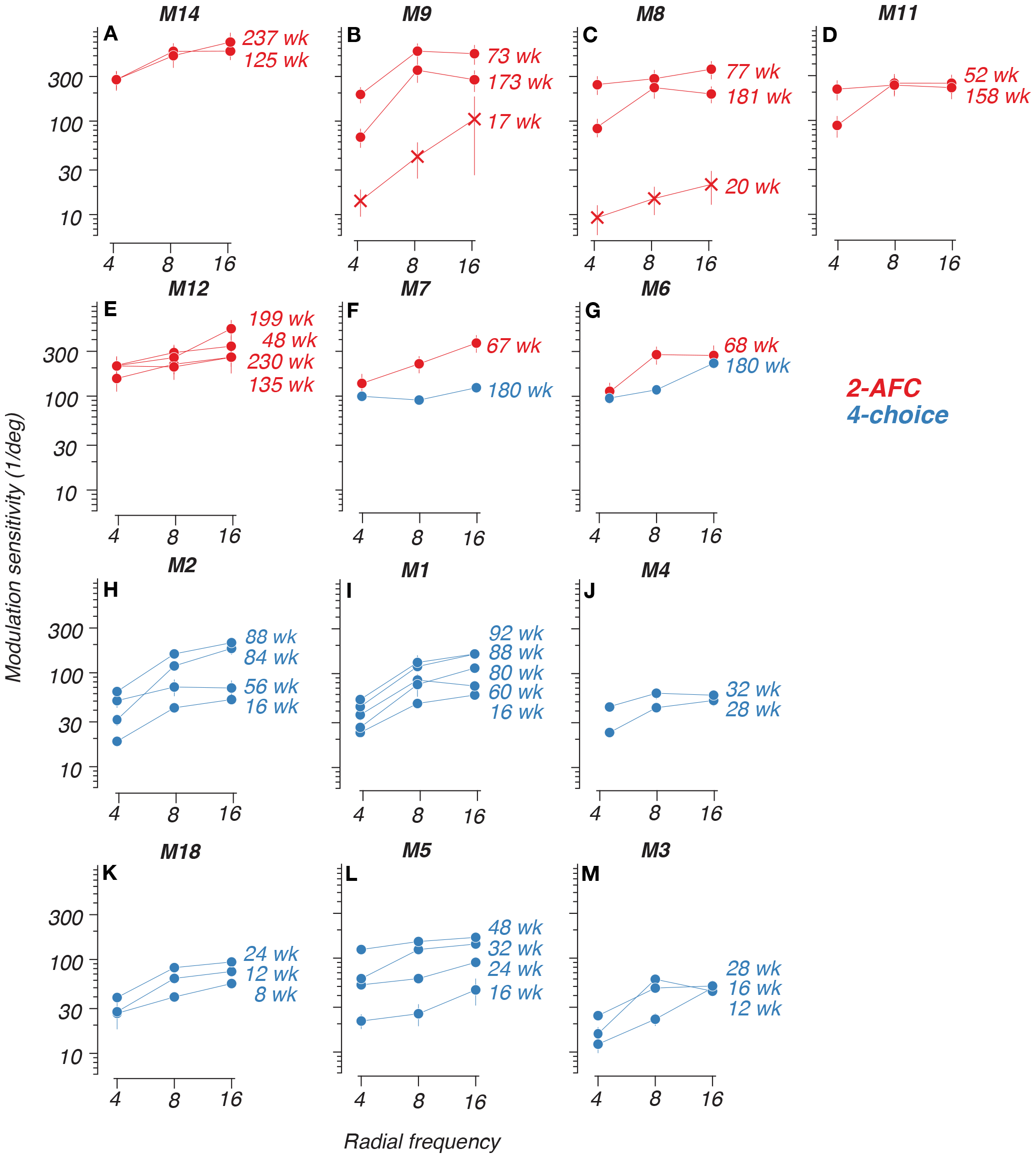
Development of sensitivity to radial frequency stimuli. Perceptual sensitivity of all subjects tested at more than one age. Sensitivity to modulation amplitude (inverse of threshold) as a function of radial frequency. Cross symbols in panels B and C correspond to reinforced-looking data. Red = 2-AFC and blue = 4-choice data. The age in weeks of the subject at the time of testing is indicated next to each curve. Error bars represent 68% CIs.

### Developmental time course of radial frequency sensitivity

To look for overall trends across all subjects and methods, longitudinal and cross-sectional data for all subjects is shown in Figure 4. The inclusion of cross-sectionally tested subjects confirms we are capturing developmental improvements rather than training effects alone, as these subjects’ sensitivities fall within the trajectory delineated by the longitudinal data. Our 2-AFC data (Figure 4 D-F) showed clear developmental asymptotes for all RFs, likely due to more data from older subjects. Meanwhile, the 4-choice data (Figure 4 A-C) had most clearly begun to asymptote for the highest radial frequencies at the older end of the age range we tested. Another interesting feature of our data is the magnitude of sensitivities computed using reinforced looking at the earliest ages tested on the 2-AFC task (cross symbols; Figure 4 D-F). These variations likely reflect the impact of temperament and engagement of the very young subjects on the difficult range of modulation amplitudes the experimenter could effectively test, yielding lower estimates of sensitivity. These data points are included in Figure 4 D-F) for completeness in visualizing our data across all subjects, but we exclude them in the analysis described below for reasons that are explained below.

**Figure 4:**
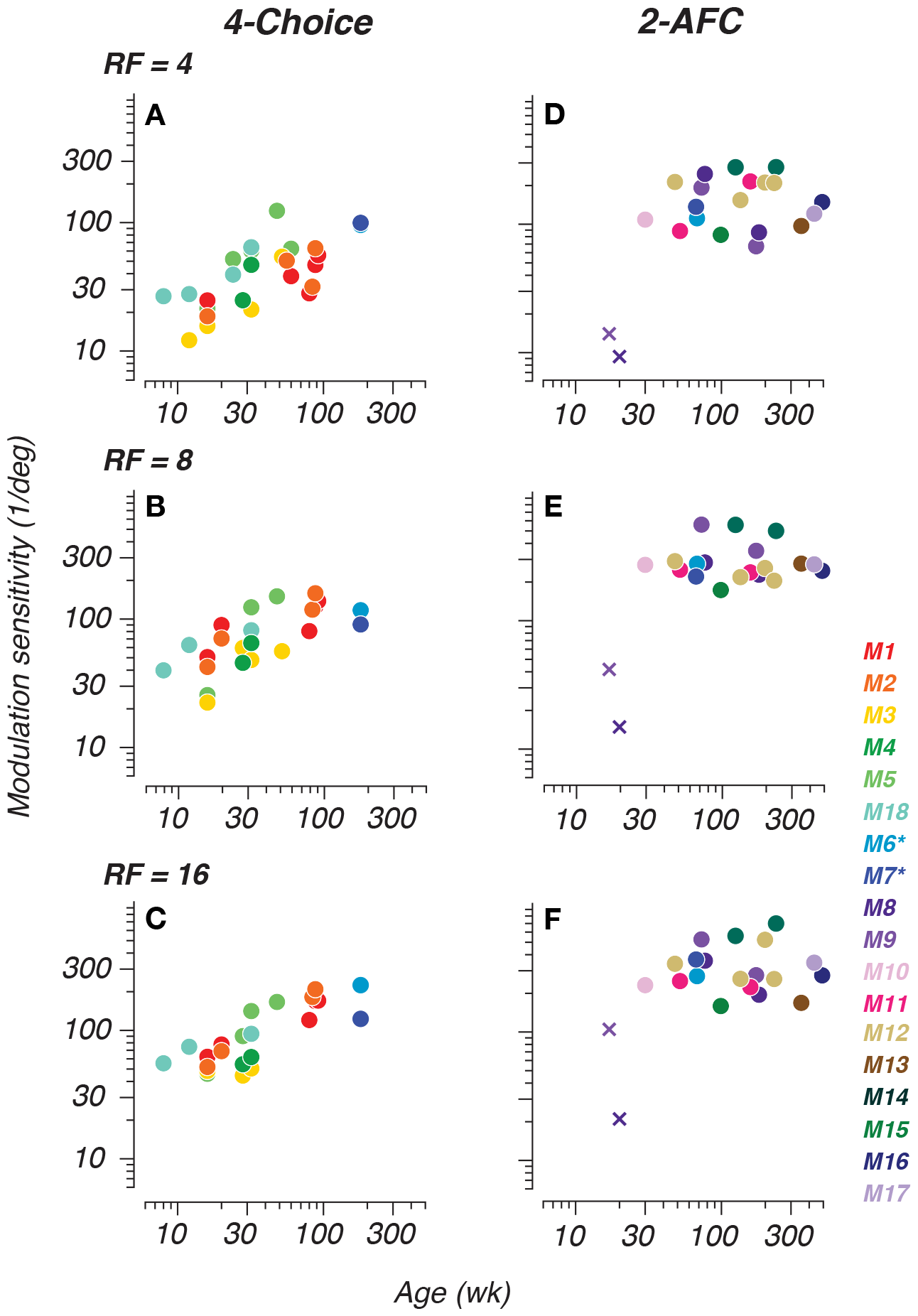
Tracking radial frequency sensitivity development with a 4-choice oddity task and a 2-alternative forced choice task. Modulation sensitivity for all subjects plotted as a function of age in weeks. Panels A-C depict data collected with the 4-choice oddity task. Panels D-F depict data collected with the 2-AFC task. Subject identity is indicated by symbol color. Cross-symbols represent reinforced-looking data. Asterisks on legend indicate that monkeys 6 and 7 were tested on both tasks.

To determine whether there was a difference between the developmental trajectories as measured with the 4-choice oddity task and traditional 2-AFC bar-pulling methods, we took account of the fact that discrimination experiments with more than one alternative increase the number of distributions of a decision variable (Kaplan, Macmillan, & Creelman, 1978). Therefore, the distributions onto which the decision variable is mapped in a 2-AFC task are not equal to those implemented by the observer on a 4-choice task and performance estimates cannot be directly compared across variable-alternative paradigms. The data from the oddity task were first adjusted by recalculating performance with a d’ = 0.95 to match performance parameters for 2-AFC tasks. We then fit Michaelis-Menten functions to the developmental trends across both populations simultaneously and for each individual subject using a constant offset fitting routine, with the semi-saturation point constrained across the two data sets *N*_2*AF C*_ = 10, *N*_4*−choice*_ = 6 and *N*_*b*_*oth* = 2. In other words, we fit the data from both tasks jointly, allowing for a constant offset to account for potential systematic differences in threshold for the two methods. We excluded the reinforced-looking data from the 2-AFC-derived set for this analysis because of the sparse number of data points collected, and this testing method may differ from either the 2-AFC and 4-choice in ways that require further examination. Exclusion of the reinforced-looking data combined with the joint fitting method allowed for more reliable estimates of the rate and extent of development. Our results show that global form perception was present by the earliest ages we tested (8-10 weeks), improved gradually, and was superior for high radial frequencies. Our results, plotted in Figure 5A-C, show that both tasks reliably capture a similar developmental trajectory for radial frequency pattern perception, with an approximately two-fold scaling difference between tasks. Peak sensitivity is represented by open circles and the fixed half-max age is represented by the open black symbol along the abscissa with horizontal/vertical bars indicating ±1 SD of the fit. Our data suggest that both types of psychophysical tasks can yield reliable performance characterizations longitudinally and cross-sectionally throughout development. Moreover, our complementary tasks provide enhanced characterization of global form perception, capturing the full developmental picture.

**Figure 5:**
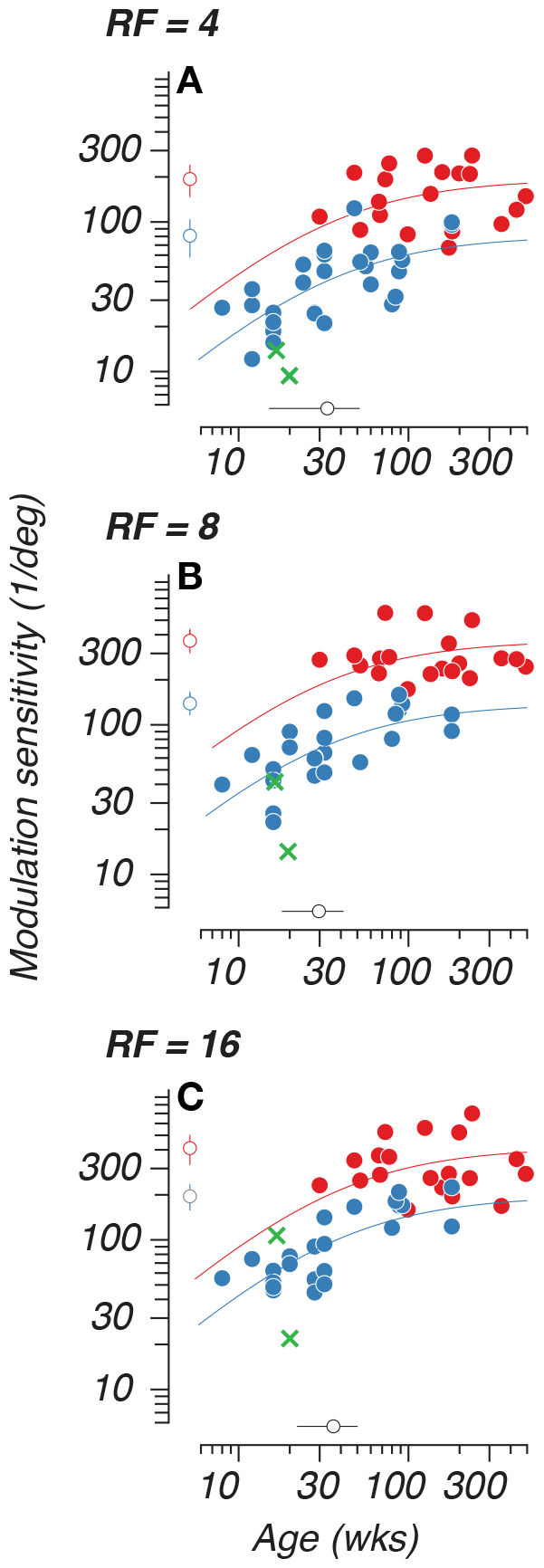
Jointly-fit 2-AFC and 4-choice sensitivity data. Panels A-C depict grouped 2-AFC and 4-choice data that were jointly fit with a constant sensitivity offset. The smooth curves represent Michaelis-Menten fits to the data across subjects for each task. The isolated open symbol intersected by vertical bars located alongside the ordinate represents the peak sensitivity value for each curve; the open symbol intersected by horizontal bars on the abscissa represents the semi-saturation age. Error bars represent ±1 SD. Green cross symbols correspond to reinforced-looking data, excluded from fitting.

Trends for the semi-saturation point and peak sensitivity across radial frequency types are plotted in Figure 6A-B. Vertical bars indicate ±1 SD of the fit. Estimated peak sensitivity was lower overall for the 4-choice oddity task at 80.95 ±23.14, 138.8 ±23.07 and 195.5 ±40 deg-1 for RF4, RF8 and RF16, respectively. On the 2-AFC task, sensitivities peaked at 193.7 ±47.08, 364.4 ±63 and 410.6 ±94.05 deg-1 for RF4, RF8 and RF16. We also found that sensitivity reached semi-saturation at 33.29 ±18.21, 29.7 ±11.72 and 36.14 ±14.05 weeks of age for RF4, RF8 and RF16 when accounting for both task modalities. Similar measures computed for contour integration and linear Glass patterns show half-max ages of 37 and 47 weeks, respectively (Kiorpes & Bassin, 2003; Kiorpes, Price, et al., 2012). The consistency between the semi-saturation point calculated here for radial frequency patterns (∼30-36 weeks) and those from previous findings helps to validate our RF methodology, and captures development as being on a similar time course as established with other global form stimuli.

**Figure 6:**
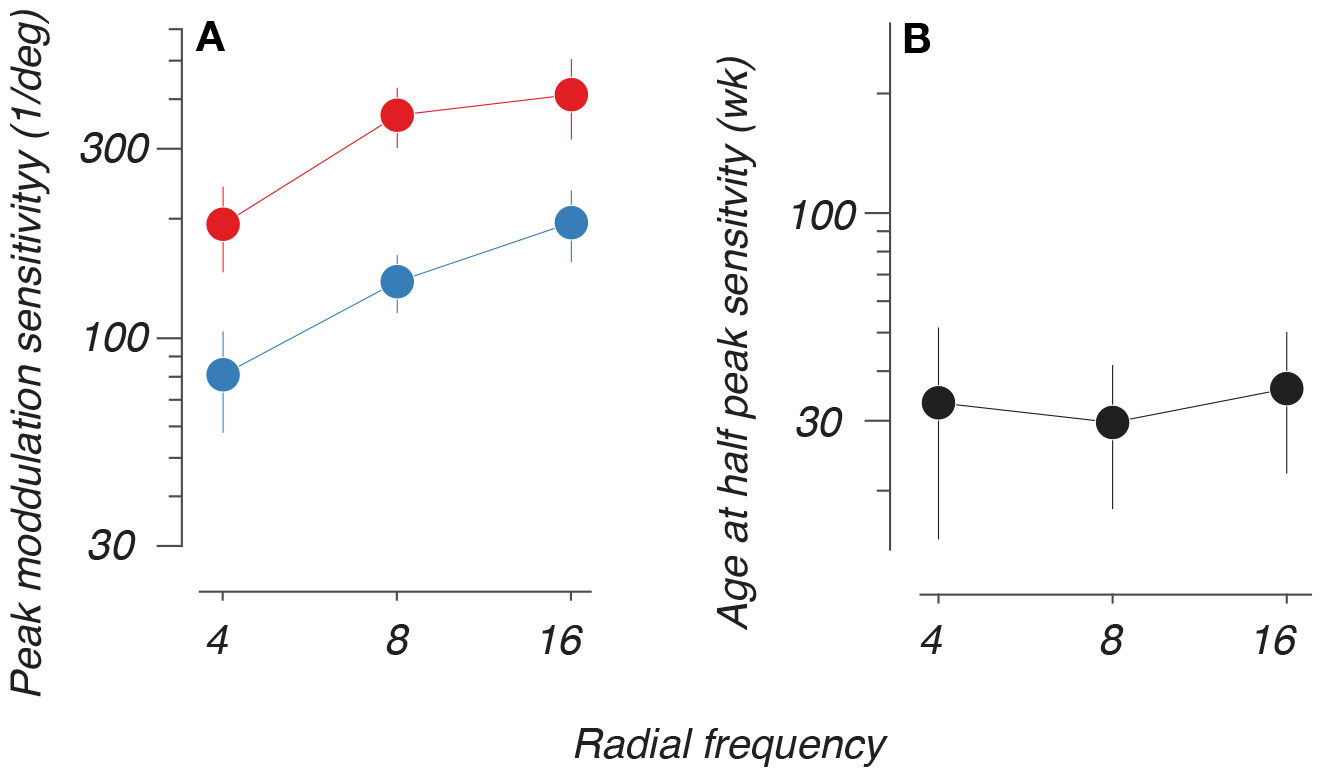
The rate and extent of the development of radial frequency sensitivity. Panel A depicts the estimated joint half-max age values for each radial frequency type. Panel B depicts the estimated peak sensitivity for both paradigms. Error bars indicate ±1 SD. Animals plotted in red were tested on 2-AFC while animals plotted in blue were tested on the 4-choice task.

We found that modulation sensitivity matures at similar rates for all three RFs tested. However, on both 2-AFC and 4-choice tasks, RF4 exhibits a significantly lower peak sensitivity compared to RF8 and RF16 (p ¡ 0.05; bootstrap hypothesis test). Modulation sensitivity for RF8 peaks lower than that for RF16, but this is not statistically significant. Despite overlap in the developmental trajectories, the observed differences in peak modulation sensitivity between low and high RFs raise the possibility for possible divergence in the form mechanisms that are selectively responsive to coarse or fine visual features.

## Discussion

In this study, we sought to characterize the developmental trajectory of infant macaques’ intermediate-level shape perception. We tested animals on a radial frequency pattern detection task in which they were required to discriminate between closed contours with radii modulated by a sinusoidal function of polar angle and contours with no modulation (circles). We used a range of radial frequencies (4, 8 and 16) and 5-6 modulation amplitudes for each to vary the difficulty of this discrimination. Our results suggest that perception of global form continues to develop well beyond the first year of life. Human studies show that sensitivity to radial frequency stimuli matures rapidly during the first year of life, but continues to improve into early childhood (Birch, Swanson, & Wang, 2000).

Our infant monkeys could discriminate all radial frequencies tested as early as 8-10 weeks postnatal, as shown using an automated 4-choice oddity task. Sensitivity for all radial frequencies improved with age, with an overall trend of higher sensitivity for higher radial frequencies. These developmental trends are also captured with separate measurements using a 2-AFC discrimination task. When compared directly, data from the two tasks improved in parallel, with a roughly 2x scaling difference between sensitivity measures using equated performance metrics.

### Time course of global form development

Circles have constant curvature along their contour, while radial frequency stimuli are created by systematically varying the frequency and degree of modulation amplitude. The visual system’s job is then to measure the variability of the curvature responses between both kinds of patterns, a computation that is not available to local curvature discrimination mechanisms (Wilkinson, Wilson, & Habak, 1998; Poirier & Wilson, 2006; Bell et al., 2007; Loffler, 2008). Our findings build on and expand prior research on the development of global form perception in infant humans and monkeys. While informative, past human work was limited by sparse longitudinal data and restricted age ranges. The current study helps address these limitations through longitudinal testing of infant macaques from 2 months to 10 years and implementation of a novel 4-choice oddity task, allowing for characterization of global form perception across development. The rate of improvement of global form perception in our study was faster initially and then slowed down over the first three years of age in primates, roughly corresponding to the first 10 years of human childhood, consistent with human research on radial frequency pattern perception (Birch, Swanson, & Wang, 2000; Wang, Morale, et al., 2009). Moreover, our findings are consistent with studies on other types of global form perception, such as contour integration and Glass pattern discrimination, which depend on spatial integration across the large regions of an image and also show protracted developmental timelines (Kiorpes & Bassin, 2003; Kiorpes & Movshon, 2003, 2004).

### Local vs. global processing

Previous studies using radial frequency stimuli have noted that thresholds are in the hyperacuity range (Wilkinson, Wilson, & Habak, 1998). Adult macaque data also reach hyperacuity range. The extended developmental trajectory for radial frequency discrimination is consistent with hyperacuity development in macaques reported by Kiorpes (1992, 2015) on the development of grating and Vernier acuity in macaques. In that study, grating acuity was relatively more mature in early postnatal life than Vernier acuity, with Vernier acuity maturing at a faster rate over a longer time course. Wilkinson, Wilson, and Habak (1998) studied the relationship between radial frequency and perceptual thresholds, finding that radial frequencies below 2 resulted in drastically higher thresholds. Radial frequencies of 3-24, including the range used in the present study, led to thresholds that plateaued between 2-9 seconds of arc. These values fall well below the cutoff for hyperacuity (Hess, Wang, & Dakin, 1999; Jeffrey, Wang, & Birch, 2002; Wang & Hess, 2005; Schmidtmann & Kingdom, 2017). The increasing peak sensitivities for higher radial frequency types observed across our subjects may therefore be due to increased information necessary for the observer to resolve spatial distinctions at scales that exceed the layout of receptors tiling the retina. Our study may also provide support for the use of radial frequency stimuli as a resource to test hyperacuity in the diagnostic process for amblyopia in children (Subramanian et al., 2012).

Jeffrey, Wang, and Birch (2002) found that radial deformation thresholds depended primarily on the circular contour frequency (cycles of modulation per degree of unmodulated contour) rather than the absolute RF. At a given radius, higher RF patterns have greater circular contour frequency. For example, at a radius of 1.5 degree: RF4 has a circular contour frequency of 0.43 cycles/degree, RF8 has 0.86 cycles/degree and RF16 has 1.72 cycles/degree. Jeffrey, Wang, and Birch (2002) showed thresholds improve linearly with log circular contour frequency until reaching a plateau around 1.3-2.6 cycles/degree. The lower peak sensitivity we observed for RF4 is likely because a circular contour frequency of 0.43 cycles/deg is well below the plateau region found by (Jeffrey, Wang, & Birch, 2002), while RF8 and RF16 at 1.5 degree radius are close to or above the plateau circular contour frequency where asymptotic sensitivity is reached. The lack of significant difference between RF8 and RF16 peak sensitivity is consistent with their thresholds both being on the plateau portion of the function. The CCF finding from Jeffrey et al. has interesting implications for the debate about whether RF perception requires spatial integration across the broader stimulus by higher visual areas. On one hand, the CCF result shows that radial frequency thresholds depend on the local spacing of modulation cycles along the contour rather than the overall stimulus size or RF number. However, the plateau in CCF tuning still extends over multiple cycles (1.3-2.6 cycles/deg), suggesting integration of local cues. Moreover, thresholds get worse if the RF segments are disconnected or phase randomized (Wilkinson, Wilson, & Habak, 1998; Hess, Wang, & Dakin, 1999; Loffler, Wilson, & Wilkinson, 2003; Jeffrey, Wang, & Birch, 2002), indicating global processing.

### 4-choice oddity and 2-AFC task comparison

Our data are novel because they provide a longitudinal characterization of global form sensitivity development using two psychophysical methods. In our comparison, we observed that both the oddity and discrimination tasks capture similar developmental trajectories with a ∼2x scaling difference in sensitivity between tasks. The difference in scaling between our data sets may reflect the inherent differences in decision variable mapping between a 4-choice oddity and a 2-AFC discrimination task, given the lower number of alternatives in the 2-AFC. Prior literature shows that oddity tasks are learned faster and more accurately than their matching or identification counterparts, given the cognitive requirement of understanding the concept of “sameness” in matching tasks (Daniel, Wright, & Katz, 2016; Miller & Shettleworth, 2007). This is to say, oddity tasks do not require understanding the stimulus content. In the oddity task, animals can saccade toward a target by evaluating *relative* aspects between the stimuli and choosing the “odd one out”. The 2-AFC discrimination, however, requires a slightly different cognitive strategy in which animals may depend on observing the *absolute* values of the relevant stimulus dimension - or its identity - to choose between a circle vs. radially deformed pattern (Bowers, 1976; Milton & Pothos, 2011). This is supported by findings suggesting that oddity discriminations activate the parietal cortex, while matching tasks are reliant on pre-frontal cortex activation (Odgaard et al., 2003; Mevorach, Humphreys, & Shalev, 2006). Practice, fatigue and/or attentional strategies may manifest differently between the two tasks and may potentially explain sensitivity differences. The 4-choice oddity task proved beneficial for collecting thousands of trials in a shorter period of time at the earlier ages we tested. Jointly modeling the complementary 2-AFC and 4-choice data sets allowed more reliable estimation of the developmental trajectory.

It is worth noting that we did notice a tendency for some animals to exhibit a bias toward one of the four possible target locations on the screen on the 4-choice oddity task. The bias varied from animal to animal and each animal’s bias shifted at different age points, which may be suggestive of shifting strategical approaches to the task requirements. Biases may evolve based on changes in reward expectations as the animals mature. However, the inclusion of catch trials, in which all stimuli were identical, allowed us to track these biases over time. Analysis of the catch trial performance revealed no consistent bias patterns across animals. While individual animals showed biases shifting between different locations at different ages, these biases were distributed randomly rather than suggesting any systematic strategic approach. Furthermore, the shifting nature of the individual animal biases indicates they were unlikely to fully account for the clear developmental improvements on non-catch oddity trials. Thus, catch trial performance suggests the observed developmental gains reflect improving perceptual abilities rather than simply emerging biases. Going forward, mapping trajectories of catch trial bias against oddity performance on a trial-by-trial basis could elucidate any potential strategic interplay and remains an open avenue for upcoming research.

### Future directions

In this study, we showed that global form perception as measured with radial frequency patterns is not fully mature in early infancy. This may reflect protracted development of visual analysis processes involved in the integration of shape signals across space. Object perception depends on a hierarchy of processing stages within the visual cortex (Van Essen, Anderson, & Felleman, 1992). A potential model of radial frequency pattern perception has been proposed by Poirier and Wilson (2006). This model sums across oriented filters to extract contour information and implements triads of oriented center-surround filters tuned to different degrees of curvature to encode local curvature signals and identify points of maximum convex curvature (Poirier & Wilson, 2006). In the last processing stage, shape is represented as curvature signal strength as a function of orientation (polar angle) around the object’s center, a computation associated with population codes in V4, which have been found to yield position invariant shape information (Pasupathy, 2006; Pasupathy & Connor, 2002). Whether postnatal maturation of V4 underlies the developmental trajectory reported here remains an open question.

### Conclusions

We found that infant macaques could report global form early in development, though this ability continues to develop well into adolescence. We find evidence of lower sensitivity of global form mechanisms to the lowest radial frequency. This likely reflects the visual system’s prioritization of high radial frequency processing critical for visual acuity and fine detail perception during development. We conclude that the neural mechanisms that integrate curvature features across space and decode increasingly smaller deviations between circles and radial frequency patterns are not fully mature by the first year of age.

## Acknowledgments

Thanks to the NYU CNS Visual Neuroscience Lab, particularly Jessica Fletcher, Tiffany Tang, Karen Fudge and Solmaz Shariat Torbaghan for training and testing our subjects. Thanks to the NYU Office of Veterinary Resources, our subjects, and the taxpaying public for funding NIH T32-MH019524 to LK, R01-EY031446 to NJM, R01-EY024914 to JAM, F31-EY031592 to CLRD and F31-EY031249 to GML.

## Notes

### Competing Interest Statement

The authors have declared no competing interest.

